# A low-cost hyperspectral scanner for natural imaging above and under water

**DOI:** 10.1101/322172

**Authors:** N. E. Nevala, T. Baden

## Abstract

Hyperspectral imaging is a widely used technology for industrial and scientific purposes, but the high cost and large size of commercial setups have made them impractical for most basic research. Here, we designed and implemented a fully open source and low-cost hyperspectral scanner based on a commercial spectrometer coupled to custom optical, mechanical and electronic components. We demonstrate our scanner’s utility for natural imaging in both terrestrial and underwater environments. Our design provides sub-nm spectral resolution between 350–1000 nm, including the UV part of the light spectrum which has been mostly absent from commercial solutions and previous natural imaging studies. By comparing the full light spectra from natural scenes to the spectral sensitivity of animals, we show how our system can be used to identify subtle variations in chromatic details detectable by different species. In addition, we have created an open access database for hyperspectral datasets collected from natural scenes in the UK and India. Together with comprehensive online build- and use-instructions, our setup provides an inexpensive and customisable solution to gather and share hyperspectral imaging data.

---

Abbreviations used. C_1–3_: Chromatic axes 1–3; MWS: Middle Wavelength Sensitive; PC: Principal Component; PCA: Principal Component Analysis; PVC: Polyvinyl Chloride; RGBU: Red-Green-Blue-Ultraviolet; SNR: Signal-to-Noise Ratio; TTL: Transistor-Transistor Logic; UHI: Underwater Hyperspectral Imager; USB: Universal Serial Bus; UV: Ultraviolet

## INTRODUCTION

Hyperspectral imaging combines spatial and detailed spectral information of a scene to construct images where the full spectrum of light at each pixel is known^1^. Commercial hyperspectral imaging technology is used, for example, in food industry^2,3^, agriculture^4,5^ and astronomy^1^. However, these devices are typically expensive, lack the ultraviolet (UV) part of the spectrum and only few work under water. Moreover, many are bulky and must be attached to a plane or other heavy machinery, which makes them unsuitable for most basic research. Here, we present a low-cost and open source hyperspectral scanner design and demonstrate its utility for studying animal colour vision in the context of the natural visual world.

Animals obtain sensory information that meets their specific needs to stay alive and to reproduce. For many animals, this requires telling wavelength independent from intensity – an ability widely referred to as colour vision. To study what chromatic contrasts are available for an animal to see in nature requires measuring the spectral content of its environment (natural imaging) and comparing this to the eye’s spectral sensitivity.

Most previous work on natural imaging to study animal colour vision used sets of spectrally narrow images generated by iteratively placing different interference filters within the range of 400–1,000 nm^6–9^ in front of a spectrally broad sensor array. So far, a major focus has been on our own trichromatic visual system that samples the short (blue “B”), medium (green “G”) and long (red “R”) wavelength (“human visible”) range of the electromagnetic spectrum^6,8,10–12^. However, across animals the number and spectral sensitivity of retinal photoreceptor types varies widely. Perhaps most importantly, and unlike humans, many animals can see in the UV part of the spectrum, which has not been included in available hyperspectral measurements from terrestrial or underwater scenes. Johnsen et al. (2013, 2016)^13,14^ used an underwater hyperspectral imager (UHI) to map the seafloor in an effort to identify structures and objects with varying depth, but more shallow underwater habitats have not been studied in this way. Finally, in 2013 Baden et al.^15^ used a hyperspectral scanner based on a spectrometer reaching the UV spectrum of light and an optical fibre controlled by two servo motors. With their setup it is possible to build hyperspectral images in a similar way to the design presented here, but the system is both bulky and fragile. In addition, their setup cannot be easily waterproofed because the point of light from the scene is guided with the optic fibre attached to the spectrometer. Our design uses mirrors instead to overcome these shortcomings.

Here, we designed and built a low-cost open source hyperspectral scanner from 3D printed parts, off-the-shelf electronic components and a commercial spectrometer that can take full spectrum (350–1,000 nm), low spatial resolution (4.7°) images above and under water. With our fully open design and instructions it is possible for researchers to build and modify their own hyperspectral scanners at substantially lower costs compared to commercial devices (~£1,500 for a spectrometer if unavailable, plus ~£113–340 for all additional components, compared to tens to hundreds of thousands for commercial alternatives). We demonstrate the performance of our system using example scans and show how this data can be used to study animal colour vision in the immediate context of their natural visual world. We provide all raw data of these and additional scans to populate a new public database of natural hyperspectral images measured in the UK and in India (https://zenodo.org/communities/hyperspectral-natural-imaging), to complement existing datasets^16–18^.

## METHODS

### Hardware design

The device is built around a trigger-enabled, commercial spectrometer (Thorlabs CCS200/M, advertised as 200–1,000 nm but effectively useful above 350 nm). A set of two movable UV reflecting mirrors (Thorlabs PFSQ10–03-F01 25.4 × 25.4 mm and PFSQ05-03-F01 12.7 × 12.7 mm) directs light from the scanned scene onto the spectrometer’s sensor region via a pinhole (see also Baden et al. 2013)^15^. To gradually assemble an image, an Arduino Uno microcontroller (www.Arduino.cc) iteratively moves the two mirrors via servo-motors along a pre-defined scan-path under serial control from a computer. At each new mirror position, the Arduino triggers the spectrometer via a transistor-transistor logic (TTL) pulse to take a single reading. An optional 9V battery powers the Arduino to relieve its universal serial bus (USB) power connection. The entire set-up is encased in a waterproofed housing fitted with a quartz-window (Thorlabs WG42012 50.8 mm UVFS Broadband Precision Window) to permit light to enter. For underwater measurements, optional diving weights can be added to control buoyancy. All internal mechanical components were designed using the freely available OpenSCAD (www.OpenScad.org) and 3D printed on an Ultimaker 2 3D printer running Cura 2.7.0 (Ultimaker). For detailed build instructions including all 3D files and Arduino control code, see the project’s GitHub page at www.github.com/BadenLab/3Dprinting_and_electronics/tree/master/Hyperspectral%20scanner.

### Scan-paths

Four scan paths are pre-programmed onto the Arduino control code: a 100 point raster at 6° x- and y-spacing (60° × 60°), and three equi-spaced spirals at *r* = ±30° at n=300, 600 or 1,000 points, respectively (Supplementary Figure 1). To generate spirals, we computed n points of a Fermat’s spiral:

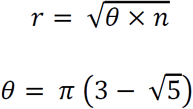
where *r* is the radius and *θ*, in radians, is the “golden angle” (~137.5°). Next, we sorted points by angle from the origin and thereafter ran a custom algorithm to minimise total path length. For this, we iteratively and randomly exchanged two scan positions and calculated total path length. Exchanges were kept if they resulted in path shortening but rejected in all other cases. Running this algorithm for 10^5^ iterations resulted in the semi-scrambled scan paths shown in SFig. 1.

### Data collection

All recordings shown in this work used the 1,000-point spiral. Acquisition time for each scan was 4–6 minutes, depending on the time set for each mirror movement (260–500 ms) and the spectrometer’s integration time (100–200 ms). These were adjusted based on the amount of light available in the environment to yield an approximately constant signal-to-noise ratio (SNR) between scans. In all cases, the scanner was supported using a hard-plastic box to maintain an upright position. All outdoor scans were taken in sunny weather with a clear sky. For details of the underwater measurement done in West Bengal India, see Zimmermann, Nevala, Yoshimatsu et al., 2017^19^. In addition, we took a 180° RGB colour photograph of each scanned scene with an action camera (Campark ACT80 3K 360°) or a ~120° photograph with an ELP megapixel Super Mini 720p USB Camera Module.

### Data analysis

All data was analysed using custom scripts written in IGOR Pro 7 (Wavemetrics) and Fiji (NIH). To visualise scanned images, we calculated the effective brightness of each individual spectrum (hereafter referred to as “pixel”) as sampled by different animals’ opsin templates. In each case, we z-normalised each channel’s output across an entire scan and mapped the resultant brightness map to 16-bit greyscale or false-colour coded maps, in each case with zero centred at 2^15^ and range to 0 and to 2^16^–1. We then mapped each pixel onto the 2D plane using a standard fish-eye projection. To map each spiral scan into a bitmap image, we scaled a blank 150×150 target vector to ±30° (same as the scanner range), mapped each of *n* scanner pixels to its nearest position in this target vector to yield *n* seed-pixels, and linearly interpolated between seed-pixels to give the final image. The 150 × 150 pixel (60 × 60 degrees) target vector was truncated beyond 30° from the centre to cut the corners which comprised no data points. We also created hyperspectral videos by adding a 3^rd^ dimension so that each pixel in the 150 × 150 target vector holds a full spectrum. This way each video is constructed from 800 individual images where one frame equals to 1 nm window starting from 200 nm.

### Principal component analysis

For principal component analysis (PCA), we always projected across the chromatic dimension (e.g. human trichromatic image would use 3 basis vectors, “red”, “green” and “blue”) after z-normalising each vector.

## RESULTS

The scanner with water-proofed casing, its inner workings and control logic are illustrated in Figure 1. Light from the to-be-imaged scene enters the box through the quartz window (Fig. 1A) and reflects off the larger and then the smaller mirror, passing through a pinhole to illuminate the active part of the spectrometer (Fig. 1B). To scan a scene, an Arduino script is started via serial command from a computer to iteratively move the two mirrors through a pre-defined scan path (Methods and Supplementary Video 1). At each scan-position, the mirrors briefly wait while the spectrometer is triggered to take a single reading. All instructions for building the scanner, including 3D part models and the microcontroller control code are provided at the project’s GitHub page at https://github.com/BadenLab/3Dprinting_and_electronics/tree/master/Hyperspectral%20scanner.

**Figure 1.**
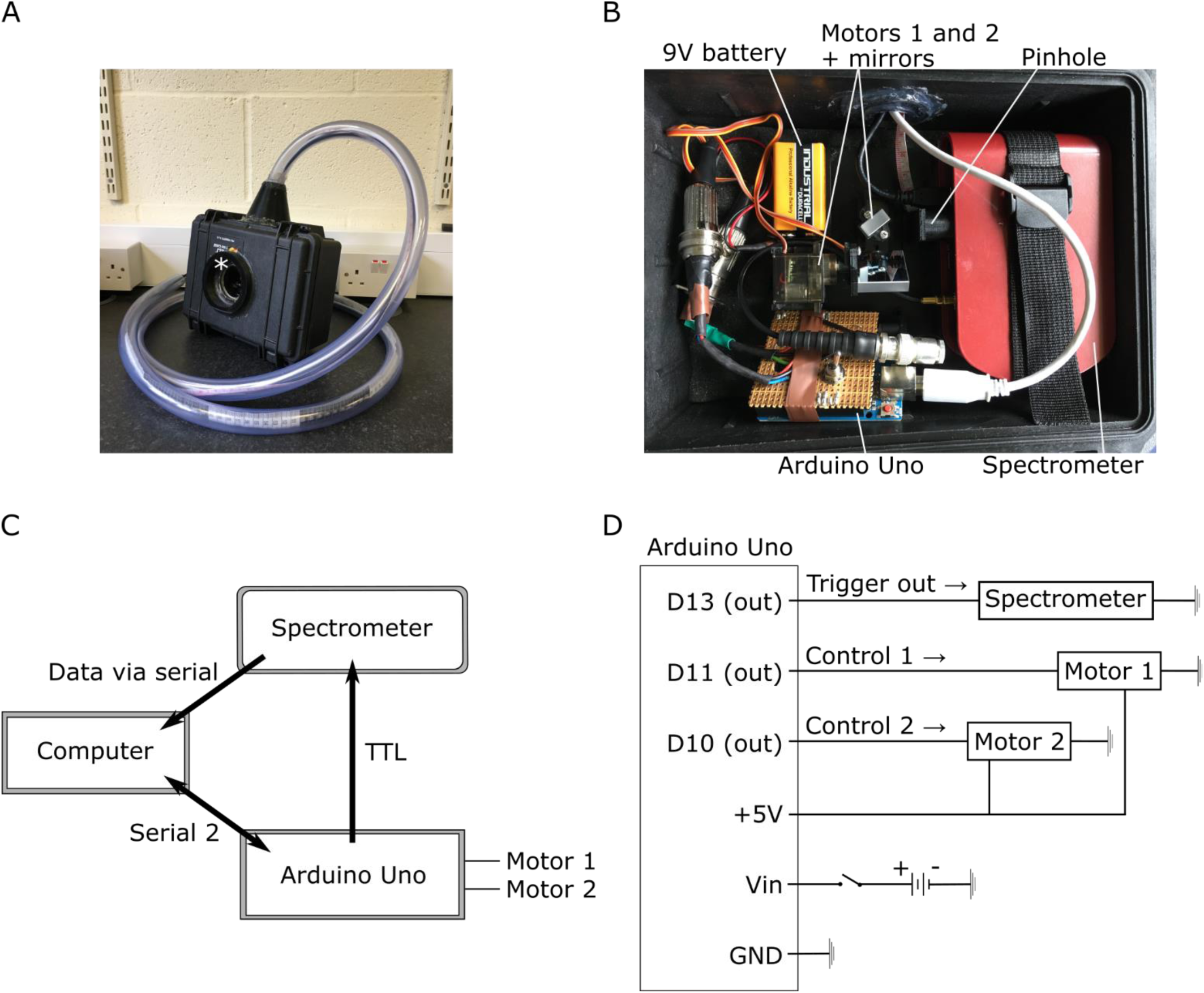
A Hyperspectral scanner for low-cost natural imaging. **(A)** The waterproof casing with a window (white asterisk) for light to enter. The PVC tube on top protects the cables to the computer. **(B)** Internal arrangement of parts: the spectrometer, Arduino Uno microcontroller, 9V battery, two servo motors (Motors 1 and 2) with mirrors attached to them and a pinhole. Light reaches first the larger mirror underneath the window of the casing, reflects to the smaller mirror and from there through the pinhole to the spectrometer’s sensor. Light deflected off the first mirror is partly shadowed by the edges of the casing, which creates dark stripes at the horizontal edges of the scanned images when the box is closed. These edges are cropped in the presented example scans (Figs. 2 and 7). Spectral filtering by the quartz window was corrected for in postprocessing (Supplementary Figure 2). **(C)** Operational logic. The scanning path is uploaded to the Arduino from the computer via Serial 2 connection to define the motor movements. After each movement the spectrophotometer is triggered via TTL to take a measurement and send the data to the computer vial serial. The ongoing state of the scanning path is fed from the control circuit to the computer. **(D)** Circuit diagram.

### Scanner performance

In our scanner design, several factors contribute to the spatial resolution limit of the complete system. These include spacing of the individual scan-points, angular precision of the servo-motors, the effective angular size of the pinhole as well as the optical properties of the mirrors and the quartz window. To therefore establish the scanner’s effective spatial resolution, we scanned a printout of 8.6° black and white bars in the mid-day sun using a 1,000-point spiral (Fig. 2A, Supplementary Figure 1) and compared the result (Methods) to the original scene (Fig. 2B, C). The difference between these two profiles approximately equates to a Gaussian blur of 2.36° standard deviation, which effectively translates to ~4.7° as the finest detail the scanner can reliably resolve under these light conditions. While this spatial resolution falls far behind even the simplest commercial digital camera systems, our scanner instead provides 650 nm spectral range and sub-nm resolution that can be used to identify fine spectral details in the scanned scene.

**Figure 2.**
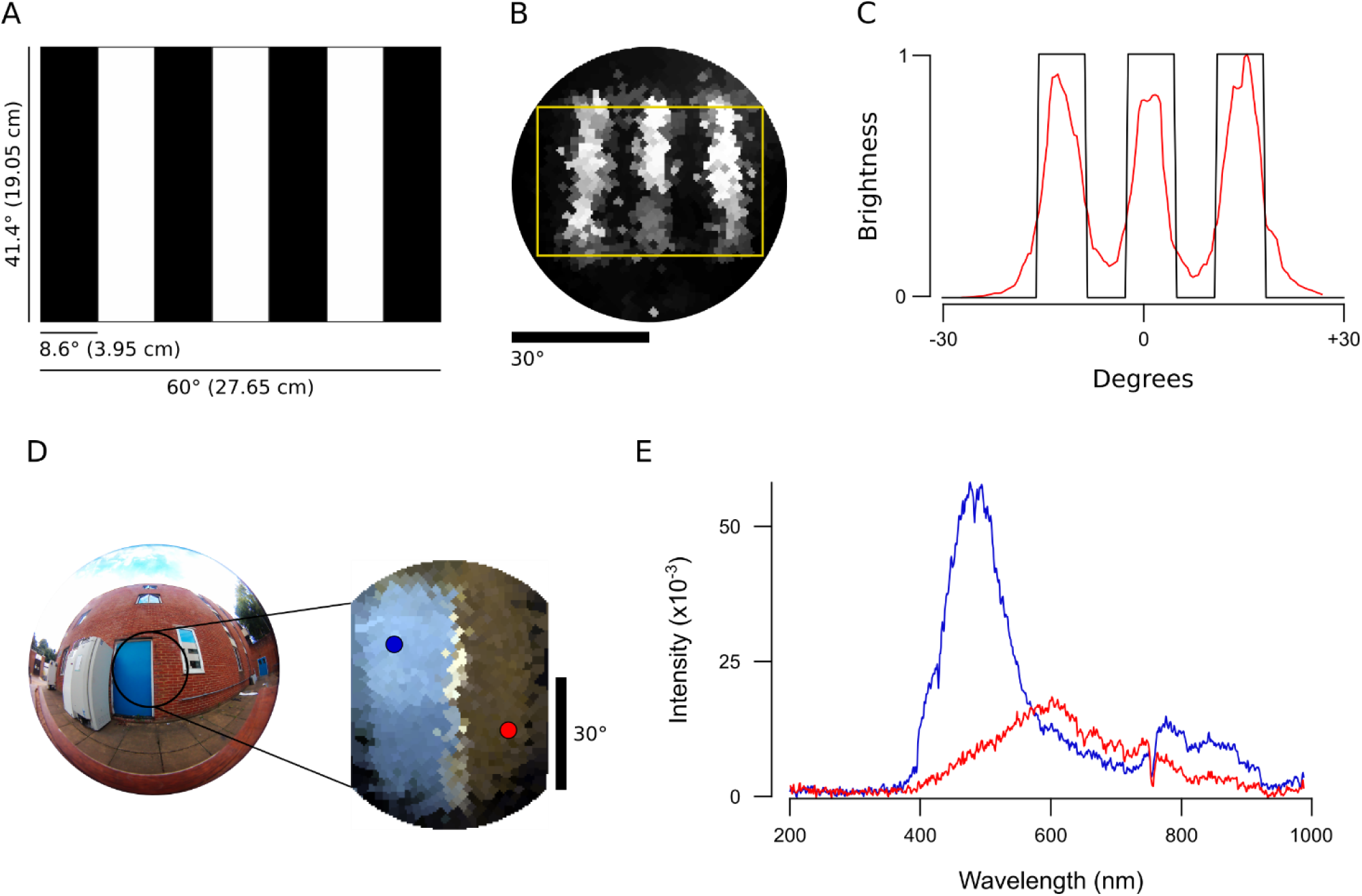
Scanner performance. (**A-C**) A printout of 8.6° black and white bars (A) was scanned with a 1,000 point spiral scanning path (B) to estimate the scanner’s spatial resolution. In (C), the average brightness (red) as indicated in (B) is plotted on top of the idealised brightness profile (black). **(D)** An action camera picture of the blue door + red brick wall measured outdoors and an RGB representation image of the scan when using opsin templates from human spectral sensitivity. Blue and red dots in the RGB representation refer to the two points used to show examples of individual spectra in **(E)**.

To illustrate the scanner’s spectral resolution, we took a 1,000-point scan in the mid-day sun of a blue door and red brick wall (Fig. 2D) and reconstructed the scene based on human red, green and blue opsin templates^20^ to assemble an RGB image (Methods, Fig. 2D). From this scan, we then picked two individual “pixels” (blue and red dots) and extracted their full spectra (Fig. 2E). Next, we illustrate the function with examples from terrestrial and underwater scenes.

### Natural imaging and animal colour vision

The ability to take high-spectral resolution images is useful for many applications, including food quality controls^2,3^, agricultural monitoring^4,5^ and surface material identification from space^1^. Another possibility is to study the spectral information available for colour vision by different animals. Here, our portable, waterproofed and low-cost hyperspectral scanner reaching into the UV range allows studying the light environment animals live in. To illustrate what can be achieved in this field, we showcase scans of three different scenes: a forest scene from Brighton, UK (Figs. 3–5), a close-up scan of a flowering cactus (Fig. 6) and an underwater river scene from West Bengal, India (Fig. 7). In each case, the estimated 60° field of view covered by the scanner is indicated in the accompanying widefield photos (Fig. 3A, 4A, 6A, 7A). To showcase chromatic contrasts available for colour vision by different animals in these scenes, we reconstructed the forest and cactus data with mouse *(Mus musculus)*, human *(Homo sapiens)*, bee *(Apis melifera)*, butterfly *(Graphium sarpedon)*, chicken *(Gallus gallus domesticus)* and zebra finch *(Taeniopygia guttata)* spectral sensitivities (Fig. 5B, 6C). The underwater scan was reconstructed based on zebrafish (*Danio rerio*) spectral sensitivity (Fig. 7B)^20–25^. In addition, we provide hyperspectral movies between 200 and 1,000 nm for these three scenes, where each frame is a 1 nm instance of the scanned scene (Supplementary Videos 2–4). These videos illustrate how different structures in the scene appear at different wavelengths.

**Figure 3.**
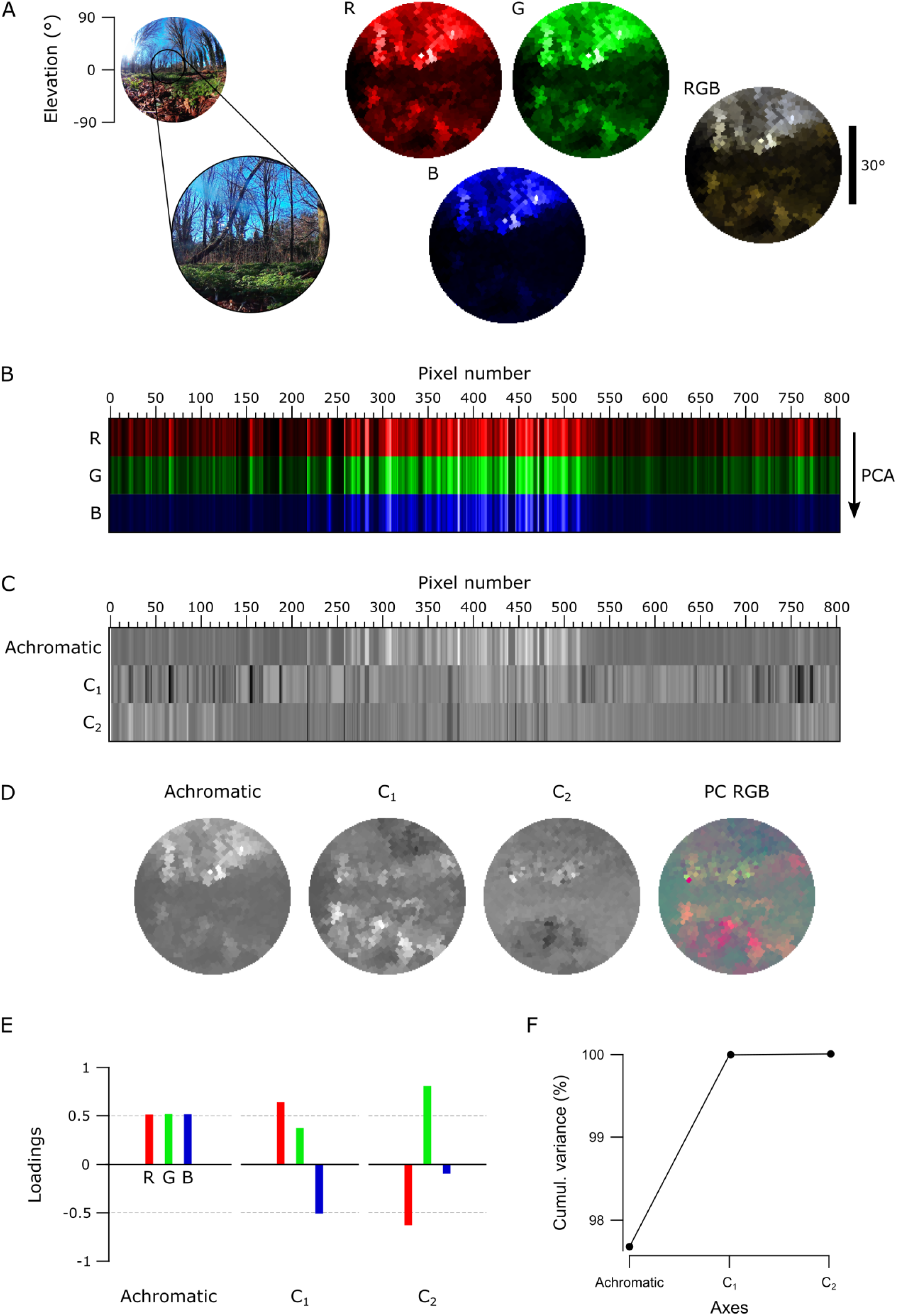
An example data set of the forest scene with human spectral sensitivity. **(A)** A 180° photo of the forest scene with an approximate 60° scanner covered area (left). On the right, monochromatic R-, G- and B-channels were constructed from the scanned data by multiplying spectra from each pixel with the opsin templates (see Fig. 5B, 6C). The RGB image shows the reconstruction built based on the opsin channels. The different colour appearance of this RGB reconstruction compared to the photograph is due to differential colour-channel equalisations in the two images. **(B)** Pixels from the R-, G- and B-channels aligned in the order of the measurement with an arrow on the right indicating the direction of the principal component analysis (PCA). **(C)** Achromatic and chromatic axes C_1–2_ aligned in the same order as in the previous image, and then reconstructed back to images in **(D)** to add the spatial information. The RGB image shows C_1_ in red and C_2_ in green (blue set to constant brightness). **(E)** Loadings from achromatic and chromatic axes, bars illustrating the amount of input from each opsin channel. **(F)** The cumulative variance explained (%) for each axis.

**Figure 4.**
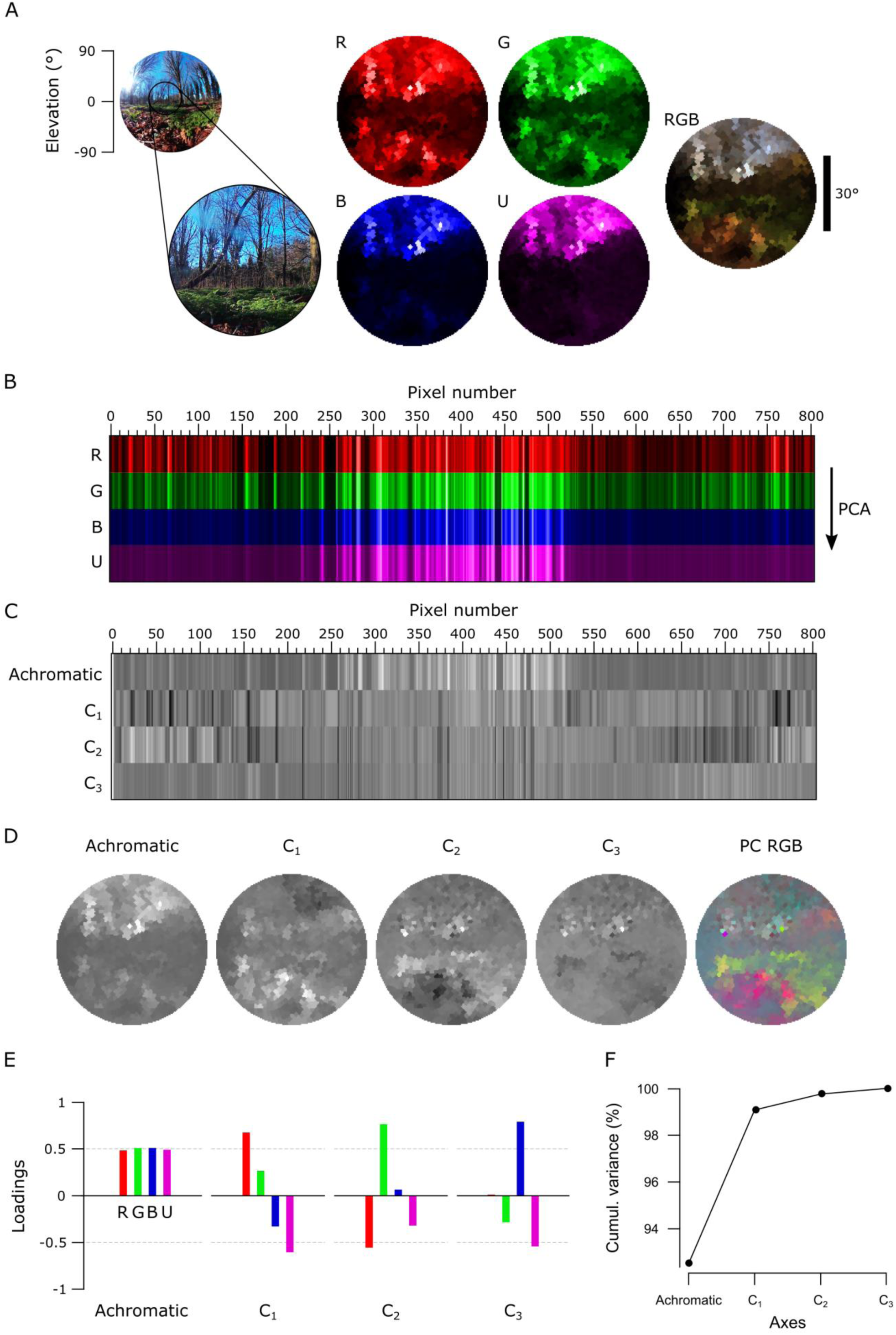
The forest scene with zebra finch spectral sensitivity. **(A)** A still image of the forest scene with the approximated 60° scanner covered area, monochromatic opsin channels (R, G, B, U) and an RGB reconstruction where R is shown as red, G as green and B+U as blue. **(B-F)** As in Fig. 3, with an addition of the UV channel (U) in all images. The RGB image in (D) displays C_1_ in red, C_2_ in green and C_3_ in blue.

**Figure 5.**
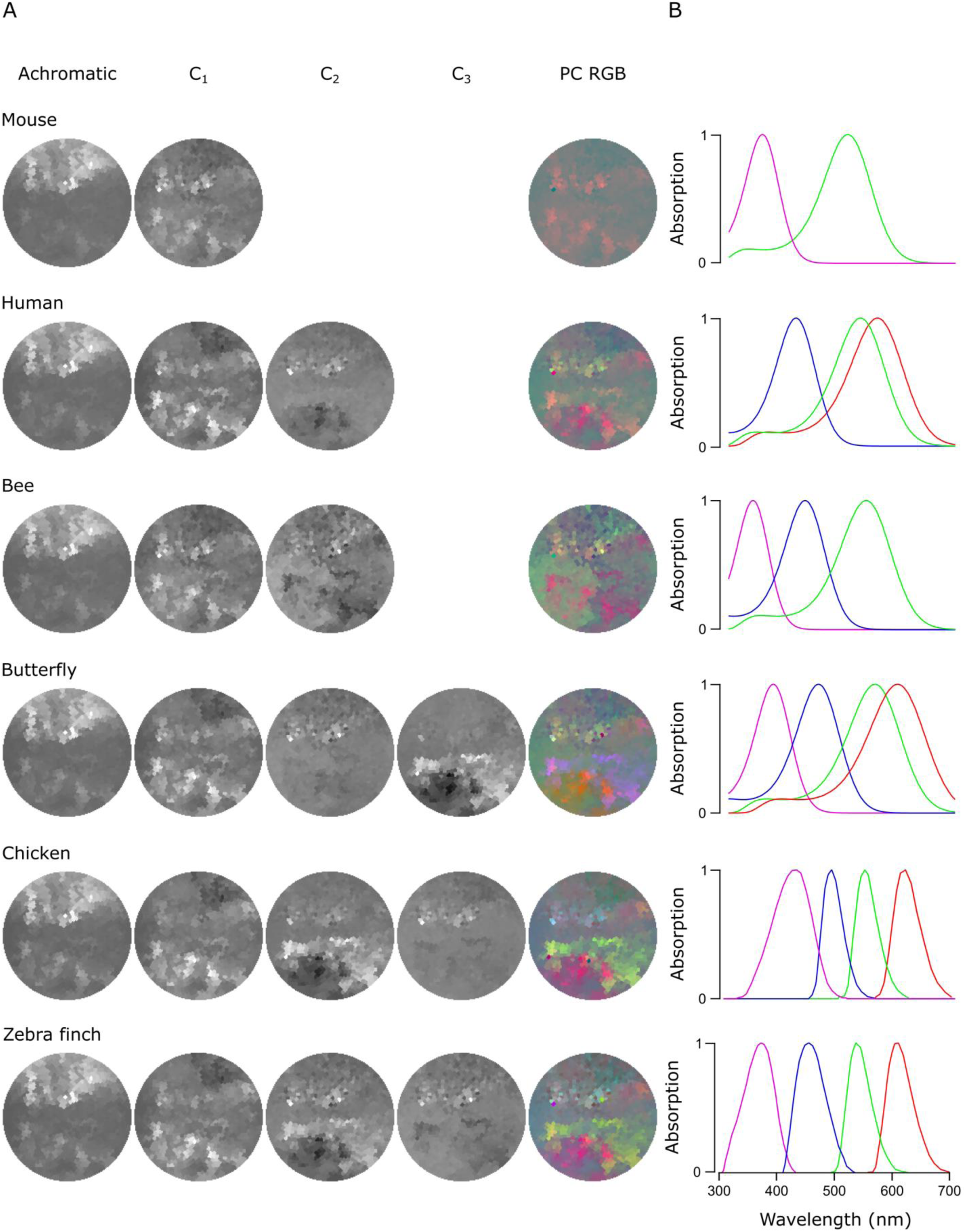
PC reconstructions of the forest scene. **(A)** Achromatic and chromatic PCA reconstructions from the forest scene data for a mouse *(Mus musculus)*, a human *(Homo sapiens)*, a bee *(Apis melifera)*, a butterfly *(Graphium sarpedon)*, a chicken *(Gallus gallus domesticus)* and a zebra finch *(Taeniopygia guttata)* and PC RGB pictures. The number of chromatic axes equals to the number of cone types minus 1. Again, the PC RGB picture is constructed from chromatic axes C_1-n_. In PC RGB, the C_1_ is shown as red, C_2_ as green and C_3_ as blue. (B) Opsin absorption curves showing the spectral sensitivity of the cones for each animal. The pink, blue, green and red curves correspond to UV, blue, green and red sensitive opsins, respectively.

**Figure 6.**
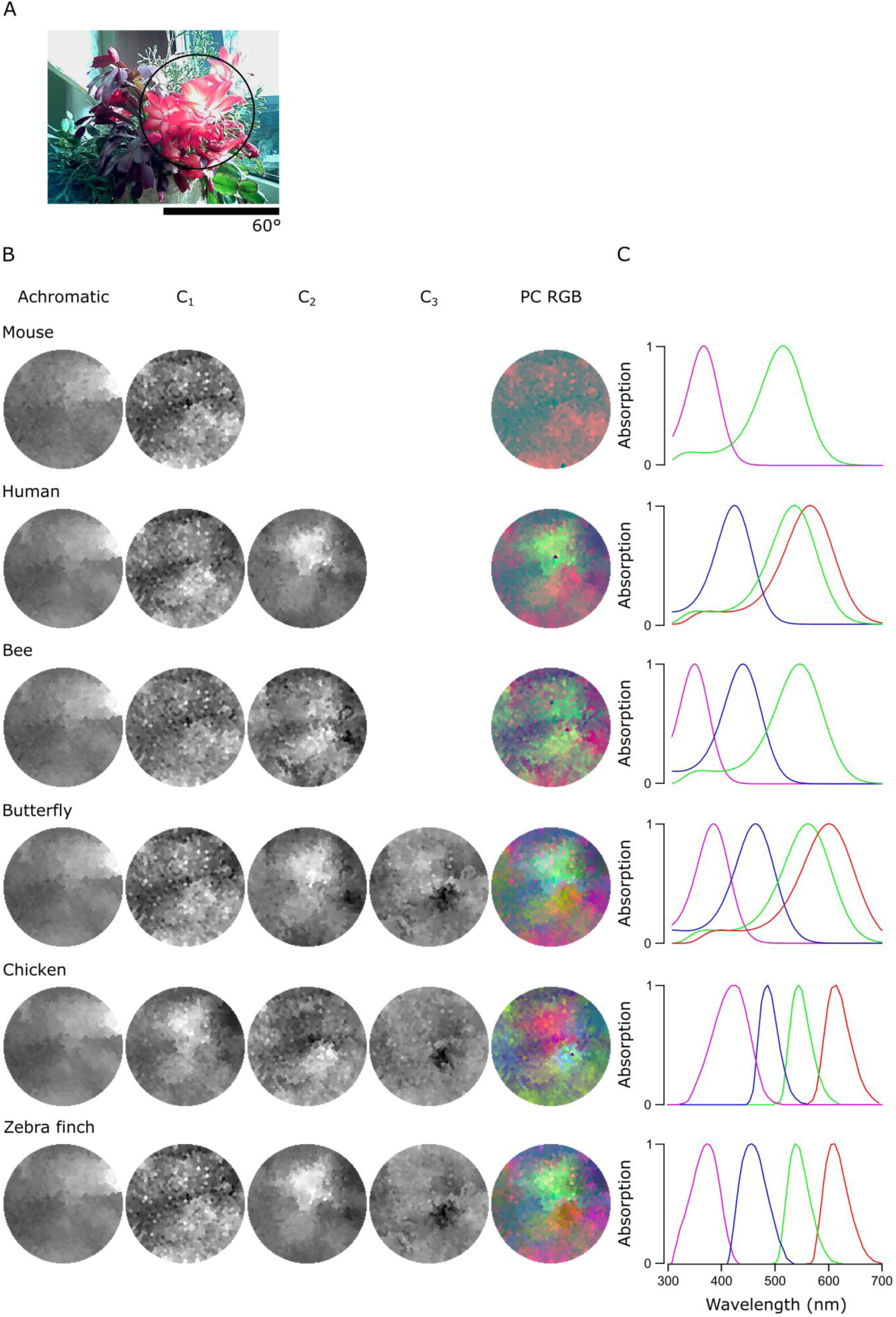
PC reconstructions of the flowering cactus. **(A)** A 120° photo of the scanned scene with a flowering cactus and the approximate 60° window (black circle) the scanner can cover. **(B)** Reconstructions for the chromatic axes C_1-n_ and PC RGB images and the absorption curves for each animal as in Fig. 5.

**Figure 7.**
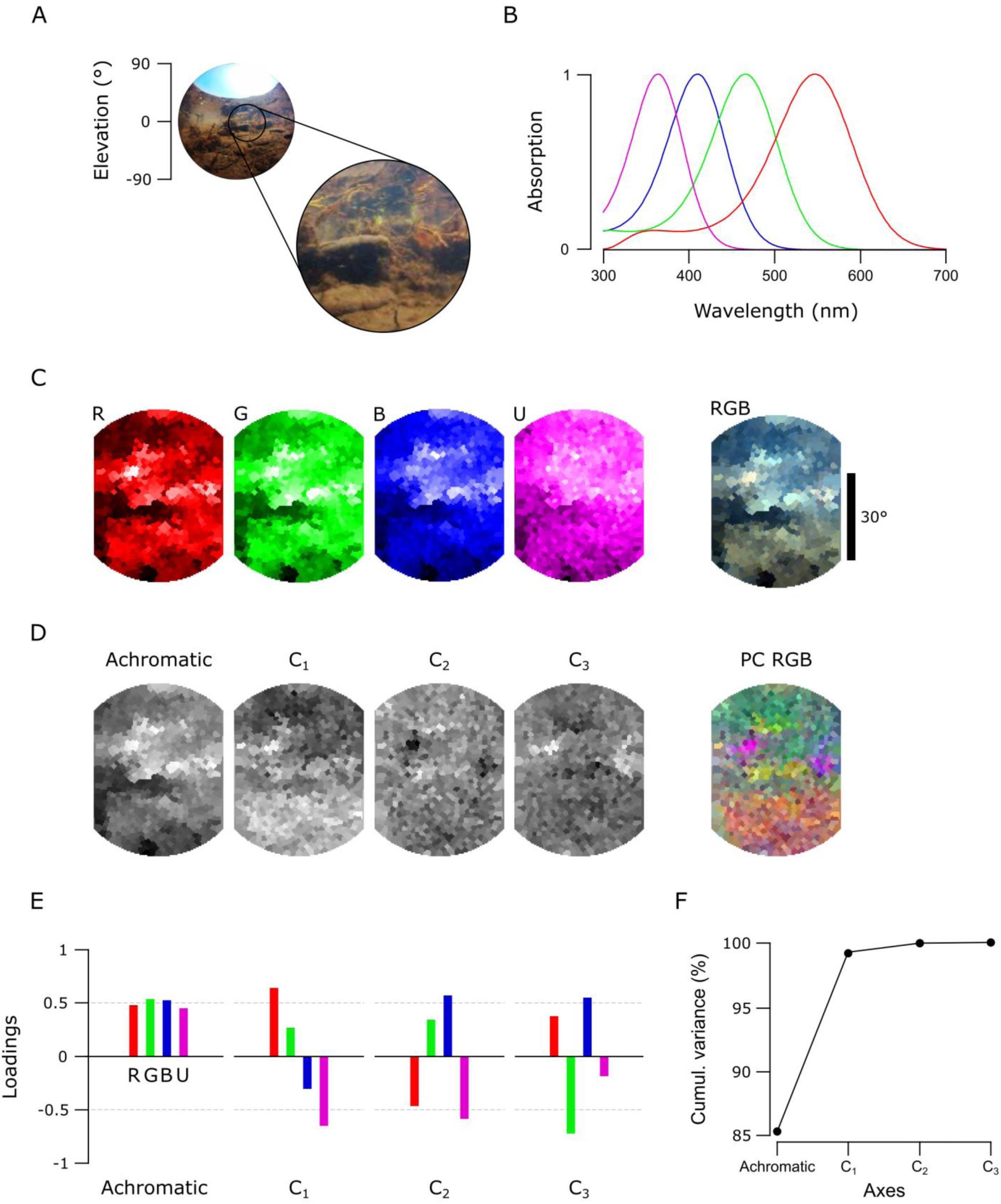
An underwater scene from India with zebrafish spectral sensitivity. **(A)** A 180° photo of the scanned underwater river scene from West Bengal, India, and the approximate 60° scanner covered window. **(B)** The zebrafish opsin complement. **(C)** The monochromatic opsin channels (RGBU) and the RGB reconstruction as in Fig. 4. **(D)** The achromatic and chromatic axes reconstructed back to images to show where in the scene information based on each axis can be found. **(E)** Loadings from each opsin channel as explained in Fig. 3E. **(F)** The cumulative variance explained (%) for each axis.

First, we used the data from the forest scene scan to compute how a trichromat human with three opsins (red, green and blue) might see it (Fig. 3). To this end, we multiplied the spectra from each “pixel” with the spectral sensitivity of each of the three corresponding opsins templates to create “opsin activation maps” (red “R”, green “G” and blue “B”, Fig. 3A, Methods), hereafter referred to as “channels”. These false-colour coded, monochromatic images show the luminance driving each opsin across the scene. In this example, the R- and G-channels clearly highlight the dark band of trees in the middle of the scene with varying light and dark structures in the sky and on the ground. However, the B-channel shows mainly structures from the sky but provides low contrast on the ground. To illustrate how these channels can be used for our sense of colour vision, we combined them into an RGB image (Fig. 3A, right).

To determine what chromatic structures are discernible with human spectral sensitivity, we used principal component analysis (PCA) across the 3-dimensional RGB space by using the R-, G- and B-channels as 3 basis vectors (Fig. 3B, C). In natural scenes, most variance across space is driven by changes in overall luminance rather than chromatic contrasts^6,9,10^. In this type of data, the first principal component (PC1) therefore reliably extracts the achromatic (greyscale) image content. From here it follows that all subsequent principal components (PC2-n) must describe the chromatic axes in the image, in decreasing order of importance. For simplicity, we hereafter refer to PC1 as the achromatic axis and PC2, PC3 and (where applicable) PC4 as first, second and third chromatic axes, respectively (C_1, 2, 3_). When applied to the example scan of the forest scene with human spectral sensitivity, the achromatic image with near equal loadings across the R-, G- and B-channels accounted for majority (97.7%) of the total image variance (Fig. 3D-F), in agreement with previous work^6,9,10^. This left 2.3% total variance for the first and second chromatic axes C_1_ and C_2_ (Table 1). In line with Ruderman et al. (1998)^6^, the chromatic contrasts emerging from PCA were R+G against B (C_1_, long- vs short-wavelength opponency) and R against G while effectively ignoring B (C_2_, Fig. 3E). These two chromatic axes predicted from the hyperspectral image matched the main chromatic comparisons performed by the human visual system (“blue vs. yellow” and “red vs. green”). To show where in the image different chromatic contrasts exist across space, and to facilitate visual comparison between animals, we also mapped the chromatic axes into an RGB image such that R displays C_1_, G C_2_ and B C_3_. Since the trichromat human can only compute two orthogonal chromatic axes (nOpsins – 1), C_3_ was set to 2^15^ (i.e. the mid-point in 16-bit) in this example. These PC-based RGB images ignore the brightness variations of the achromatic channel, therefore describing only chromatic information in a scene. This specific projection allows a trichromat human observer viewing an RGB-enabled screen or printout to judge where in a scanned scene an animal might detect dominant chromatic contrasts, even if that animal uses more than three spectral cone types for colour vision. The power of this approach can be illustrated when considering non-human colour vision based on the same dataset.

**Table 1.** The total variance explained by chromatic axes C_1-n_ in the forest and cactus scans.

| Variance explained by chromatic axes C<sub>1-n</sub> (%) |  |  |
| --- | --- | --- |
|  | Forest (Fig. 5) | Cactus (Fig. 6) |
| Mouse | 2.6 | 8.0 |
| Human | 2.3 | 1.4 |
| Bee | 3.9 | 8.1 |
| Butterfly | 3.8 | 3.8 |
| Chicken | 6.7 | 2.9 |
| Zebra finch | 7.5 | 6.5 |

Unlike humans, many animals use the ultraviolet (UV) part of the spectrum for vision^26,27^. To illustrate how the addition of UV-channel can change available chromatic information, we next performed the same analysis for a tetrachromatic zebra finch (Fig. 4). This bird uses four, approximately equi-spaced opsins (red, green, blue and UV), which in addition are spectrally sharpened with oil droplets^23^. As before, the monochromatic opsin-channels (RGB and “U” for UV, Fig. 4A) appeared with R- and G-channels showing structures both in the sky and on the ground while B- and U-channels mainly highlighted the sky. We next computed the principal components across the now four opsin channels (Fig. 4B-F).

This time the achromatic axis explained only 92.5% of the total variance leaving 7.5% for chromatic comparisons, which now comprised three chromatic axes (C_1–3_, Table 1). As with humans, the most important chromatic axis compared long- and short-wavelength channels (C_1_, R+G against B+U, single zero crossing in Fig. 4E). C_2_ was also similar to the human version by comparing R- and G-channels, but in addition paired the R-channel with the UV and the G-channel with the blue (two zero crossings). While the spatial structure highlighted by C_1_ was similar to that of the human, C_2_ picked up additional details from the ground (Fig. 4D). Finally, C_3_ (R+B against G+U) highlighted additional structures in the scene that are largely invisible to the human observer.

### An animal’s opsin complement dictates discernible chromatic contrasts

To further survey how an animal’s opsin complement can affect the way chromatic details are detectable in complex scenes, we compared data from the forest scene (Fig. 5) to a close-up scan of a flowering cactus (Fig. 6) and filtered each using different animals’ spectral sensitivities: a dichromat mouse, a trichromat human and bee and a tetrachromat butterfly, chicken and zebra finch. In these scenes, the order of the chromatic axes was largely stable across opsin complements used (PC1 – achromatic, C_1_ – long vs short wavelengths, C_2_ – R+U vs G+B, C_3_ – R+B vs G+U), and here we only show the achromatic and C_1–3_ reconstructions alongside the PC RGB images (Fig. 5A and 6B) next to the spectral sensitivity of each animal (Fig. 5B and 6C). In each case, the number of chromatic channels shown corresponds to the number of an animal’s cone types minus 1.

The chromatic axes usable by different animals revealed diverse spatio-chromatic structures from both scenes (Fig. 5 and 6). Across all animals compared, while C_1_ still reliably highlighted a long- vs. short-wavelength axis, the exact image content picked up along C_1-n_ varied between opsin complements (Fig. 5A and 6B). For example, in the cactus scene the C_1_ for the chicken highlighted spatial structures in the image that other animals instead picked up with C_2_. A similar difference was also seen in the forest scene, where C_2_ and C_3_ in butterfly showed structures that were captured in the inverse order in the chicken and zebra finch (Fig. 5A). In addition, humans and butterflies had more consistent arrangement and structures in chromatic axes between each other than with other animals, possibly due to their similarly overlapping spectral sensitivities of the green and red cones.

For all animals in both scenes, the achromatic image content captured at least 91.9% of the total variance, leaving 1.4–8.1% for the chromatic axes (Table 1). For the forest scene, the addition of opsin-channels increased the amount of variance explained by the chromatic axes, and in particular for animals with widely spaced spectral channels (e.g. with chicken and butterfly, Table 1). In general, more chromatic details was discerned with more cones, especially when these cones had low-overlap spectral sensitivities covering a wide range of the natural light spectrum (e.g. from around 350 nm to over 600 nm as with zebra finch). Moreover, spectral sharpening of the opsin peaks through the addition of oil droplets (chicken and zebra finch) brought out further details and higher chromatic contrasts in the scanned scene. The order of importance for the chromatic axes that optimally decompose scans depended strongly on the set of input vectors – the spectral shape and position of the animal’s opsins.

### Hyperspectral imaging under water

As light travels through the water column, water and dissolved particles absorb both extremes of the light spectrum making it more monochromatic with increasing depth^9,28^. Mainly because of this filtering and scattering, underwater light environments have spectral characteristics that differ strongly from terrestrial scenes. To illustrate one example from this underwater world, we show a scan from a shallow freshwater river scene (Fig. 7A) taken in the natural habitat of zebrafish *(Danio rerio)* in West Bengal, India^19^. The data was analysed based on the spectral sensitivity of the tetrachromatic zebrafish with red, green, blue and UV sensitive cones (Fig. 7B)^21,26^. In this example, the monochromatic R-, G-, and B-channels picked up different dominant spatial structures in the scene, while the U channel appeared more “blurry” with only small intensity differences around the horizon (Fig. 7C). Here, the total variance explained by the chromatic axes C_1–3_ (14.7%, Fig. 7F) was higher compared to the two terrestrial scenes. C_1_ compared long (R+G) and short (B+U) wavelengths between upper and lower parts of the scene (Fig. 7D, E) that arose from spectral filtering under water. Finally, C_2_ and C_3_ brought out further details that probably correspond to pieces of the imaged vegetation.

### An open database for natural imaging

Based on these and other additional scans above and under water from around the world (for example, see Zimmermann et al., 2017^19^) we created an open access database online (https://zenodo.org/communities/hyperspectral-natural-imaging). All measurements in the database are taken with the hyperspectral scanner as described here.

## DISCUSSION

We have designed and implemented an inexpensive and easy-to-build alternative to commercial hyperspectral scanners suited for field work above and under water. Without the spectrometer (~£1,500), the entire system can be built for ~£113–340, making it notably cheaper than commercial alternatives. In principle, any trigger-enabled spectrometer can be used for the design. Alternatively, spectrometers can also be home-built^29,30^ to further reduce costs.

The spatial resolution of the scanner with the 1,000-points scan (~4.7°), though substantially below that of most commercial camera systems, is close to the behavioural resolution limit of several model-animals like zebrafish larvae (~3°)^31^ or fruit flies (~1–4°)^32^. Notably, most animal visual systems inherently combine a low-spatial resolution chromatic representation of the visual world with a high-spatial resolution achromatic representation^33–35^. As such, our system can likely also give useful insights into the chromatic visual world of animals with much more highly resolved eyes. The spatial resolution of our system could principally be further improved, for example by using a smaller pinhole in combination with higher-angular-precision motors. However, the amount of natural light for vision is limited, especially when imaging under water where light is quickly attenuated with increasing depth. As a result, higher spatial resolution in our system would require a substantially increased integration times for each pixel. This would result in very long scan-durations, which is unfavourable when scanning in quickly changing natural environments.

Spatial resolution aside, the spectral range and detail of our scanning approach far exceeds the spectral performance of interference filter-based approaches, as used in most previous hyperspectral imaging studies^6,8,9,17,36^. This difference may be crucial for some questions. For example, zebrafish have four opsin-genes for middle wavelength sensitive (MWS) cones (“green cones”) that are used in different parts of the retina and are separated in spectral sensitivity by few nanometres^22,37^. Most interference filter setups use relatively broad spectral sensitivity steps and would therefore miss small details in the natural scenes that could be picked up with slightly different spectral sensitivities of different opsins. By choosing individual “pixels” and the spectra they hold, it is possible to analyse fine details in complex scenes that animals can use for colour vision. This can be done already with very coarse spatial resolution to reveal structures that otherwise would remain undetected. In agreement with previous studies, we have shown how principal component analysis aids to separate achromatic and chromatic information in natural images^6,9^. Here, PCA across the chromatic channels highlights spatio-chromatic aspects in the scene that may be useful for vision. Perhaps not surprisingly, this reveals major, overall trends in landscapes (Figs. 3–5) with short wavelength dominated sky and long wavelength dominated ground. This is true also for the underwater habitats (Fig. 7), where light spectrum in the water column transforms from “blue-ish” short wavelength dominated to “red-ish” long wavelength dominated with increasing depth^19^. The PCs can also highlight details in complex scenes that might otherwise stay hidden but that may be important for animals to see in their natural habitats.

## CONCLUSION

We have shown how our simple, self-made scanner can produce hyperspectral images that can be used to study animal colour vision. We have also started to populate an open database of hyperspectral images from various natural scenes (https://zenodo.org/communities/hyperspectral-natural-imaging). In the future, it will be interesting to survey a more varied set of habitats and, for example, to compare how closely related animal species living in different habitats have evolved with varying visual abilities. This could also include variations of the presented design, for example to scan larger fields of view, or a time-automation mode by which the same scene can be conveniently followed over the course of a day. We will be pleased to facilitate other’s additions to the design through a centralised project repository (www.github.com/BadenLab/3Dprinting_and_electronics/tree/master/Hyperspectral%20scanner) and hope that in this way more researchers will be able to contribute to building a more global picture of the natural light available for animal vision on earth.

---

**Supplementary Figure 1.**
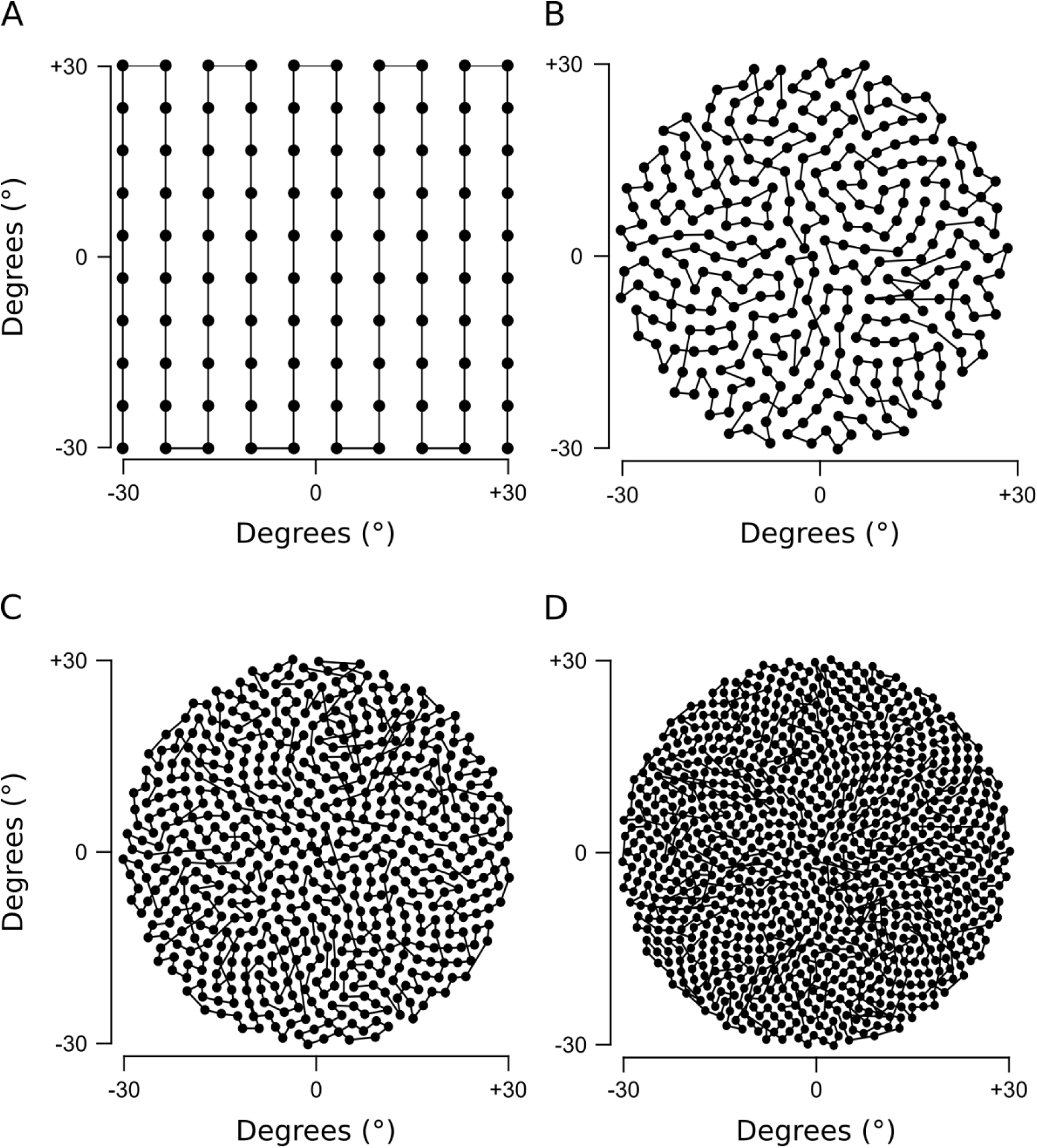
Four scanning paths created with the Fermat’s spiral across the 60° area. **(A)** 100 points square, **(B)** 300 points spiral **(C)** 600 points spiral **(D)** 1000 points spiral.

**Supplementary Figure 2.**
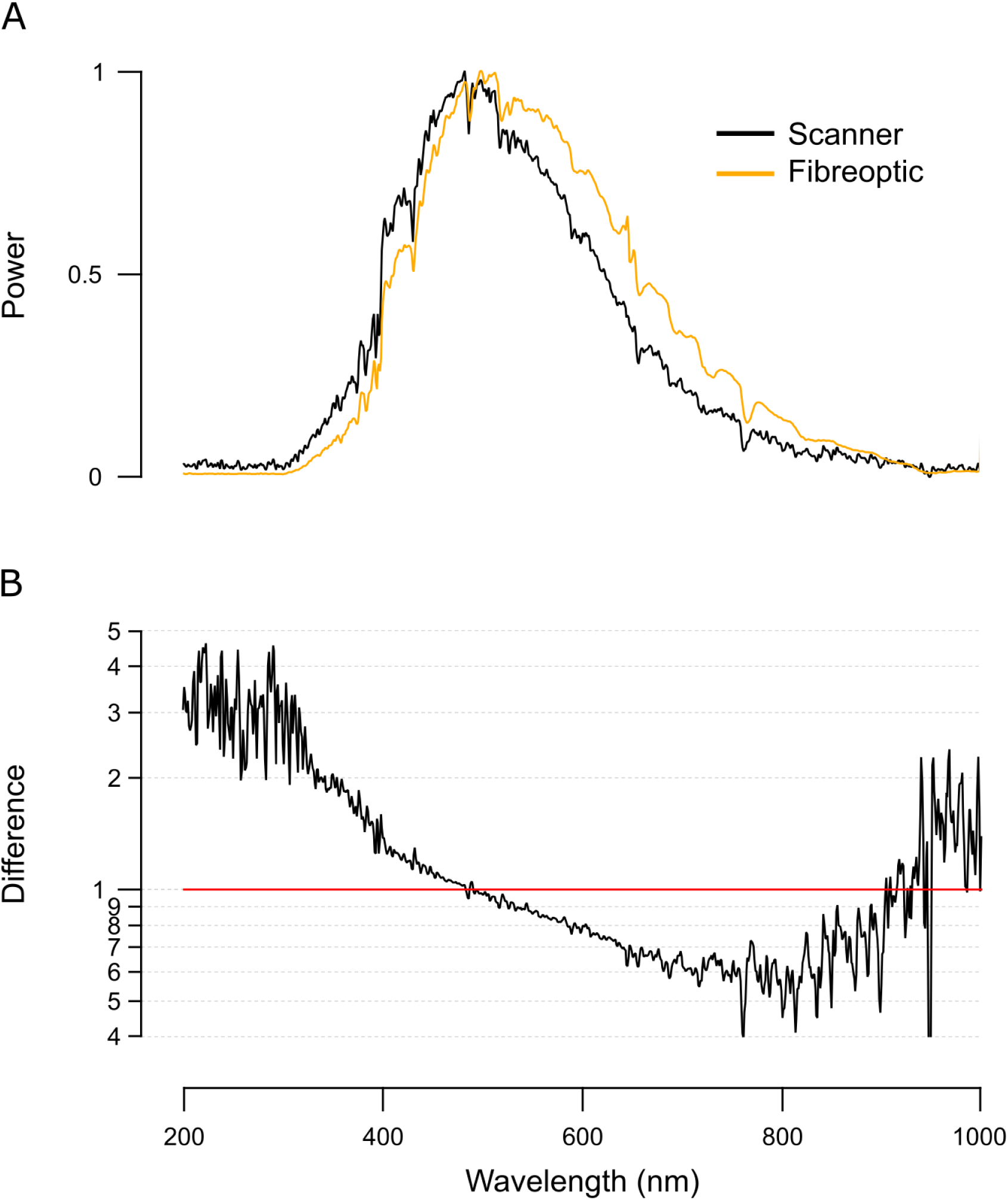
Light spectrum with and without the box. **(A)** Spectrometer readings of a clear daylight sky taken through the spectrometer’s fibreoptic (orange) or through the complete optical path of the scanner (black, i.e. 2 mirrors and a quartz window, though lacking the fibreoptic). When purchased, the spectrometer is calibrated with the fibreoptic attached. Accordingly, we computed the corresponding correction curve and applied it to all scanner data presented throughout this work **(B)**.

**Supplementary Video 1**.

A video demonstrating the mirror movements and how light is guided to the spectrometer through them.

**Supplementary Videos 2–4**.

Hyperspectral reconstructions of the three scanned scenes presented in this work, with each frame corresponding to a 1 nm instance.

## Author contributions

The scanner was conceived and implemented by NEN and TB. Data was analysed by NEN using custom scripts written by TB and modified by NEN. The paper was written by NEN with help from TB.

## Acknowledgements

We thank Kripan Sarkar and Fredrik Jutfeld for help with fieldwork, and Dan-Eric Nilsson and Daniel Osorio and Thomas Euler for general discussions. The authors would also like to acknowledge support from the FENS-Kavli Network of Excellence. Funding was provided by the European Research Council (ERC-StG “NeuroVisEco” 677687 to TB), The Medical Research Council (TB, MC_PC_15071) and the Leverhulme Trust (PLP-2017–005 to TB).

